# A systematic investigation on the involvement of complement pathway in diabetic retinopathy

**DOI:** 10.1101/656314

**Authors:** Shahna Shahulhameed, Sushma Vishwakarma, Jay Chhablani, Mudit Tyagi, Rajeev R Pappuru, Subhabrata Chakrabarti, Inderjeet Kaur

**Affiliations:** Prof Brien Holden Eye Research Centre, LV Prasad Eye Institute, Hyderabad, India; Smt. Kanuri Santhamma Center for Vitreo Retinal Diseases, LV Prasad Eye Institute, Hyderabad, India

## Abstract

**Background:** Complement system play a crucial role in retinal homeostasis. Several proteomic studies have shown deposition of complement protein in ocular tissues from diabetic retinopathy, however, their exact involvement in pathogenesis of DR remains unclear.

**Methods:** We evaluated major complement pathway proteins in the classical and alternative pathway including C1q, C4b, C3, CFB and CFH in vitreous humor and serum samples from PDR patients and controls by western blotting. Quantitative real time (QRT) PCR was done for PDR, NPDR and no-DM controls for correlating the expression of several key pro and anti -angiogenic genes with their correspondingprotein levels. Inflammation in the vitreous humor samples was assessed by ELISA and metalloproteinase activity measured by gelatin zymography. Glial activation and its association with complement activation in diabetic eyes was assessed by immunohistochemistry.

**Results:** A significant increase in C3 proteins, its activated fragment C3bα’ (110kDa) along with a concurrent up regulation of CFH was observed for PDR vitreous. QRT identified a significant upregulation of angiogenic genes and downregulation of antiangiogenic genes in PDR and NPDR cases. PDR vitreous had increased MMP9 activity and upregulation of inflammatory markers IL8, sPECAM and down regulation of anti-inflammatory marker IL-10. Increased C3 deposition and CFH upregulation were observed in DM retina. CFH was found co-localizing with CD11b+ve activated microglial cells in inner nuclear layer of DM retina.

**Conclusions:** The present study confirms increased activation of alternative complement pathway in PDR. The co-localization of CFH in CD11b +ve cells further suggests microglia as a source of CFH in diabetic retina. Increased CFH levels could be a feedback mechanism for arresting excessive complement activation DR eyes.

## Introduction

Retina being an immune privileged organ, has its own unique immune regulatory mechanisms including retinal neurons and RPE, and immune defense mechanisms comprising microglial population and the complement system. The retinal immune defense mechanism get alerted with any kind of noxious signals and starts a series of inflammatory events as an adaptive response to restore the homeostatic balance[1]. Low-level activation of the innate immune mechanisms, specifically complement system is required for preserving normal eye homeostasis and to maintain the retinal integrity while aging[2]. However, these protective mechanisms can cause detrimental consequences if the insults persist for a longer duration and lead to irreversible functional loss as evident from various neurodegenerative diseases such as Alzheimer’s, Parkinson’s, Amyotrophic lateral sclerosis, Age related macular degeneration etc[3].

Beyond its role as an immune defense mechanism, studies have shown the involvement of complement system in various tissue remodeling process such as liver regeneration, synaptic pruning while development and also in retinal angiogenesis[4-6]. The role of complement in angiogenesis have prime importance since there are several blinding eye diseases associated with abnormal ocular angiogenesis such as retinopathy of prematurity (ROP), age related macular degeneration (AMD), proliferative diabetic retinopathy (PDR) etc[7]. Both inhibitory as well as promoting role of complement are known in various ocular angiogenic conditions. A study done by Bora *et al,* in 2005, identified C3 and MAC complex deposits in neovessels in mice model of laser induced choroidal neovascularization (CNV), while the C3 knock out (C3-/-) CNV mice showed an absence of neovessels formation with reduced level of angiogenic factors thereby suggesting complement component C3 as a proangiogenic factor[8]. On the contrary, Langer *et al* in a mice model of ROP, had shown anti-angiogenic property of complement system where C3 and C5aR were required for inhibiting the polarization of macrophage towards its angiogenic potential[6]. Our earlier study (2017) on ROP identified microglia mediated excessive complement activation in vitreous of ROP babies when compared to that of age matched control babies, further suggesting complement as a promoting factor for ocular angiogenesis[9]. In past, genetic studies done in AMD had shown a strong association of *CFB* and *CFH* gene polymorphism with disease pathogenesis and strong deposition of complement components in the RPE-Bruch’s layer[10-12]. The balance between angiogenic and anti-angiogenic factors determine the extent of neovascularization[13]. *VEGF* is a potent angiogenic factor, whereas *THBS1* is both angiogenic inhibitor and a potent activator of *TGF*β. *TGF*β activation is also required to maintain the RPE mediated immune privilege status of retina[14].

DR is an important ocular angiogenic condition, that displays association with complement component deposits in retina and vitreous. It is one of the most devastating causes of irreversible vision loss with complex pathophysiology and has a global prevalence of 34.6% [15]. Gerl *et al* in 2002, for the first time identified extensive complement C3d and MAC complex (C5b-9) deposits in diabetic retinal choriocapillaries by immunohistochemistry [16]. Later, another study identified the deposition of C5b-9 complex and reduction of glycosylphosphotidyl inositol anchored inhibitors of complements such as CD55 and CD59 in the walls of retinal vessels of diabetic eyes, suggesting for the involvement of alternative complement pathway in diabetic eyes[17]. Additionally, an association of increased systemic level of iC3b/C3 and Bb/Fb was identified with the microvascular abnormalities in diabetic eyes [18]. Several vitreous proteomicsreports have shown the presence of complement proteins such as C3, CFI, CFB, C4A, C4B, C2, C4BPA, CFD, CFH in PDR subjects however the data on the levels of complement components across different studies is highly variable and does not explain clearly how they contribute to DR pathology[19-22]. Likewise, only few studies have observed the associations of polymorphisms in complement pathway genes such as *C5 (rs17611), CFH (rs800292)* and *CFB (rs1048709)* with DR[23, 24].

Thus, while the involvement of complement pathway in the DR pathogenesis was suggested based on the presence of complement proteins in the vitreous, a clear understanding on how complements could contribute to DR pathology is lacking till date. A systematic validation and comparison of alternative and classical complement pathway proteins might address the lacunae and further the knowledge on their role in PDR pathogenesis. The present study, attempted a systematic evaluation and validation of alternative and classical complement pathway proteins in PDR pathogenesis in an extended cohort. In addition, we have also correlated our findings with the expression of complement proteins in retinal tissues obtained from diabetic cadaveric donors and blood samples of DR patients and controls. Thus, the mRNA expression of pro and anti-angiogenic genes and their correlation with complement C3 and CFH expression was compared among patients and controls. Next, we correlated the level of complement activation with inflammation in PDR vitreous by analyzing inflammatory and proinflammatory cytokines by ELISA. Our study identified, elevation of C3 in PDR vitreous, especially 110kDa C3bα’ fragment and a concurrent upregulation of CFH in PDR vitreous. To the best of our knowledge, this is the first report on upregulation of CFH levels in PDR vitreous as revealed through western blotting. Most importantly, increased CFH levels only in the vitreous and not in systemic circulation further, strengthened its major role in the pathogenesis of DR. Additionally, we were also able to show that the increased complement activation correlated with the presence of activated microglia in diabetic retina.

## Materials and methods

### Enrolment of study participants and sample preparation

The study was performed according to the guidelines of Declaration of Helsinki and approved by Institutional Review Board. Vitreous samples (100 µl) were collected from control (n=100) and PDR subjects (n=100) while undergoing pars planar vitrectomy after obtaining written informed consent. Samples were collected in surgery rooms under aseptic conditions and then immediately transferred to the lab in ice cold condition. The samples were then centrifuged at 14,000 rpm for 10 min at 4°C to remove any cellular debris and then stored at −80 degrees till further use. Proteins were lysed in equal volume of RIPA buffer and precipitated with ice-cold acetone overnight at −80°C. The precipitated proteins were collected by centrifugation at 14,000 rpm for 1hour at 4°C and dissolved the protein pellet in 1x PBS containing protease inhibitor. Blood samples were collected in a vacutainer without the anticoagulant from PDR (n=38), NPDR (n=38) and control (n=38) subjects and separated the serum within 1 hour of sample collection by centrifugation at 1500g for 15 minutes and then stored at −80°C. The total protein concentration was calculated by bicinchoninic acid (BCA) assay. The detailed demographics of the vitreous and serum samples used in this study is given in the supplementary table1 and 2.

### Western blotting

Western blotting was performed in vitreous and serum samples for identifying the role of complement pathway in PDR pathogenesis. Total C3 and its fragmentation pattern in serum samples were compared among PDR, NPDR and no-DM subjects. The quantitative expression of classical complement pathway proteins C1q and C4b was evaluated in vitreous humor of PDR and no-DM subjects. Alternative complement pathway proteins C3, CFB and CFH were evaluated both in serum and in vitreous. Standard protocol for western blotting were followed[9]. The details of antibodies used and their dilutions are given in supplementary table 3. The blots were developed and protein specific bands were visualized in a LICOR image scanner using LI-COR image studio software and the band intensities were quantified.

### Immunohistochemistry (IHC)

Cadaveric control (n=3) and diabetic eyes from type 2 DM with no retinopathy (n=3) were collected in a sterile moist chamber within 24 hours of death from Ramayamma international eye bank, LV Prasad Eye institute, Hyderabad, India, according to the tenets of Declaration of Helsinki. The retinas were removed carefully from the eyes under a dissection microscope and fixed them in 4% formalin and paraffin sections were made. For IHC, antigen retrieval was done for the deparaffinized tissue sections using Tris citrate buffer of pH 6. The sections were permeabilized using methanol for 30 min at −20°C followed by washing thrice with 1X PBS. Blocking was done with 2% BSA followed by sections were incubated with primary antibodies for overnight at 4°C (Ms C3-Santacruz, 1:50, sc-28294, Ms CFH-Santacruz, sc-166613, 1: 50, Rb CXCR4, sc-9036, 1:50, Rb CD11b, CST,49420, 1:200, Rb GFAP, Dako, Z0334). After washing thrice with 1x PBS and the sections were incubated with appropriate fluorescent labelled secondary antibodies (Goat anti Rb594,Life Tech. A-11012, 1:300, Goat anti Ms 594, Life Tech. A-11005, 1:300, Goat anti Rb 488, Life Tech. A-11008, 1:300) for 1hour at room temperature. The sections were counterstained with DAPI and visualized the staining under fluorescent microscope (EVOS) using appropriate filters.

### Gelatin zymography

Gelatin zymography was done to analyze the matrix metalloproteinases activity (MMP2 and MMP9) in the vitreous samples obtained from PDR and control subjectsas per the protocol [25] using 10 µg of vitreous proteins.

### Enzyme Linked Immunosorbent Assay

ELISA was done to evaluate the level of cytokines such as sPECAM, IL-8 and IL-10 in the vitreous samples collected from PDR and control subjects following the standard protocol [9]. Vitreous humor samples were diluted with assay buffer to 1:3 dilution. Quantitative data was obtained using Luminex system with xPONENT® software and the generated results were exported in terms of median fluorescent intensity and calculated the concentration of the analytes in the PDR and controls. The significance was calculated based on the t-test with a *p* value <0.05.

### RNA isolation and quantitative real time PCR

Blood samples were collected in K3EDTA coated 3mL blood vacutainers from PDR, NPDR and no-DM subjects and RNA was isolated using Trizol-chloroform method. 1µl of RNA was converted into cDNA using iScript cDNA conversion kit (1708891, Bio-Rad) as per manufacture’s protocol. Semi-quantitative PCR were performed on 7900 HT system using TaqMan assay chemistry for *C3, TGF*-β and SyBr green chemistry for *VEGF, CFH* and *CXCR4*. β-actin was used as normalization control using standard thermal cycling conditions. Cycle threshold (CT) values for each test gene were obtained for each sample using the *SDS2.3* software and fold change was calculated using 2^-ΔΔCt^. The primer sequence used for qRT are given in the supplementary table 4.

## Results

### 1. Systematic evaluation of complement pathway activation by analyzing the central complement protein C3

Complement component C3, is the central complement proteins which converge all the three pathways of complement system. Our vitreous proteomic analysis (manuscript under submission) identified several peptides for complement pathway proteins in the vitreous samples of DR patients (see the supplementary table 5). To study further, western blotting was done for C3 in vitreous samples obtained from PDR and no-DM controls, level of total C3 and its activated fragments were evaluated (Fig1a). A significant increase in total C3 was observed in vitreous PDR (1.9±0.25, *p*\*<*0.004*) (n=38) compared to no-DM controls (0.98±0.18) (n=38) (Fig1b). The activated C3 fragments such as intact C3 consists of 195kDa in size, 110 kDa size C3bα’, C3α (120kDa),C3β of 75kDa, α-1 fragment of iC3b (65kDa) and C3c α’ fragment-2, (43kDa) were observed in PDR and no-DM controls. A uniform pattern of C3 fragmentation was not observed in vitreous samples, therefore fragments of C3 in each of the samples were analysed and compared. A significant upregulation in C3bα’of 110kDa fragment was seen in PDR vitreous compared to controls (Fig1c). While our western blotting of C3 in PDR, NPDR serum samples identified a slight increase in total C3 compared with no-DM serum however, this increase was not statistically significant (PDR vs no-DM: 1.69±0.58, *p*>*0.05,* NPDR vs no-DM: 1.38±0.24,*p*>*0.05* and PDR vs NPDR 1.19±0.23, *p*>*0.05*). Additionally, we did not identify significant changes in any of the C3 fragments in serum samples.

### 2. Contribution of classical pathway of complement activation in diabetic retinopathy

The classical pathway of complement activation was evaluated in vitreous and serum samples by western blotting using classical pathway proteins such as C1q and C4b (Fig 2a and 2c). The levels of C1q and C4b proteins in the vitreous were not significantly different across PDR and controls (C1q-PDR: 1.81±0.44, controls: 1.32±0.38 *p*>*0.05,* C4b-control: PDR: 1.39±0.4, 1.08±0.2, *p*>*0.05*) (fig 2c and 2d). Likewise, no significant change of C1q level was observed in serum samples of PDR and NPDR as compared to no-DM controls (PDR: 1.81±0.44, controls: 1.32±0.38 *p*>*0.05*). Thus, the classical pathway of complement activation was not involved for complement activation in DR.

**Fig 1a.**
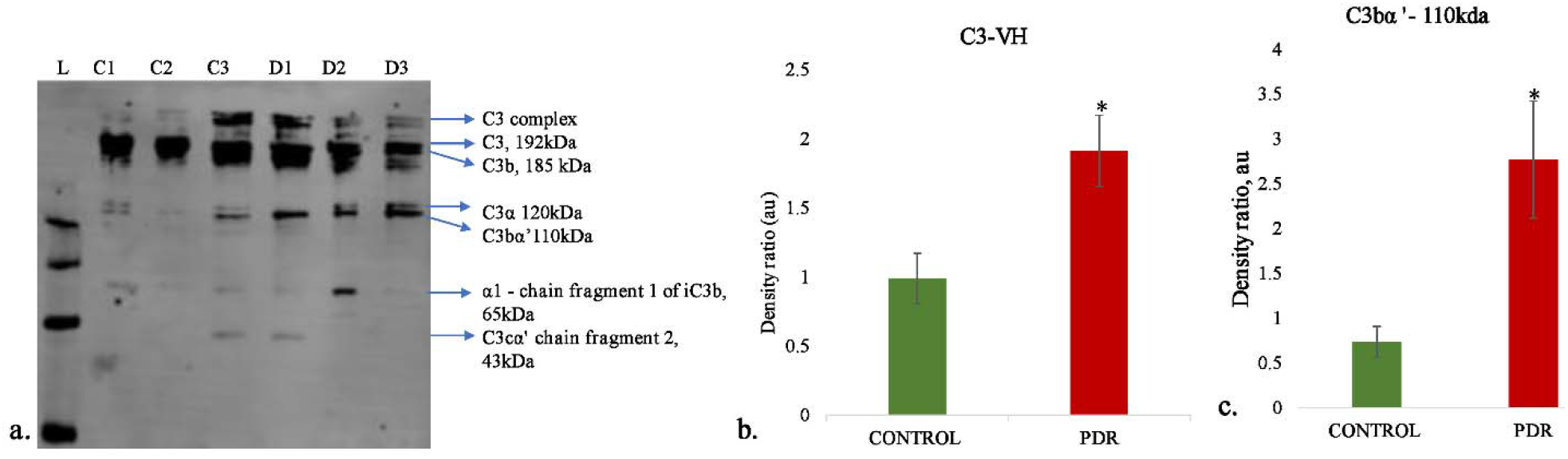
Representative western blot of C3 in PDR no-DM vitreous (b). Quantification of total C3 in PDR and in no-DM control vitreous by densitometry (PDR, n=38 and Control, n=38), Quantification of C3bα’ (110 kDa, in PDR (n=13) vitreous compared to control vitreous (n=13) *P*<0.006*, \**p<0.05,* Data represented as Mean ±SEM, C-control vitreous, D-PDR vitreous, L-protein ladder, VH-Vitreous Humor.

**Fig 2.**
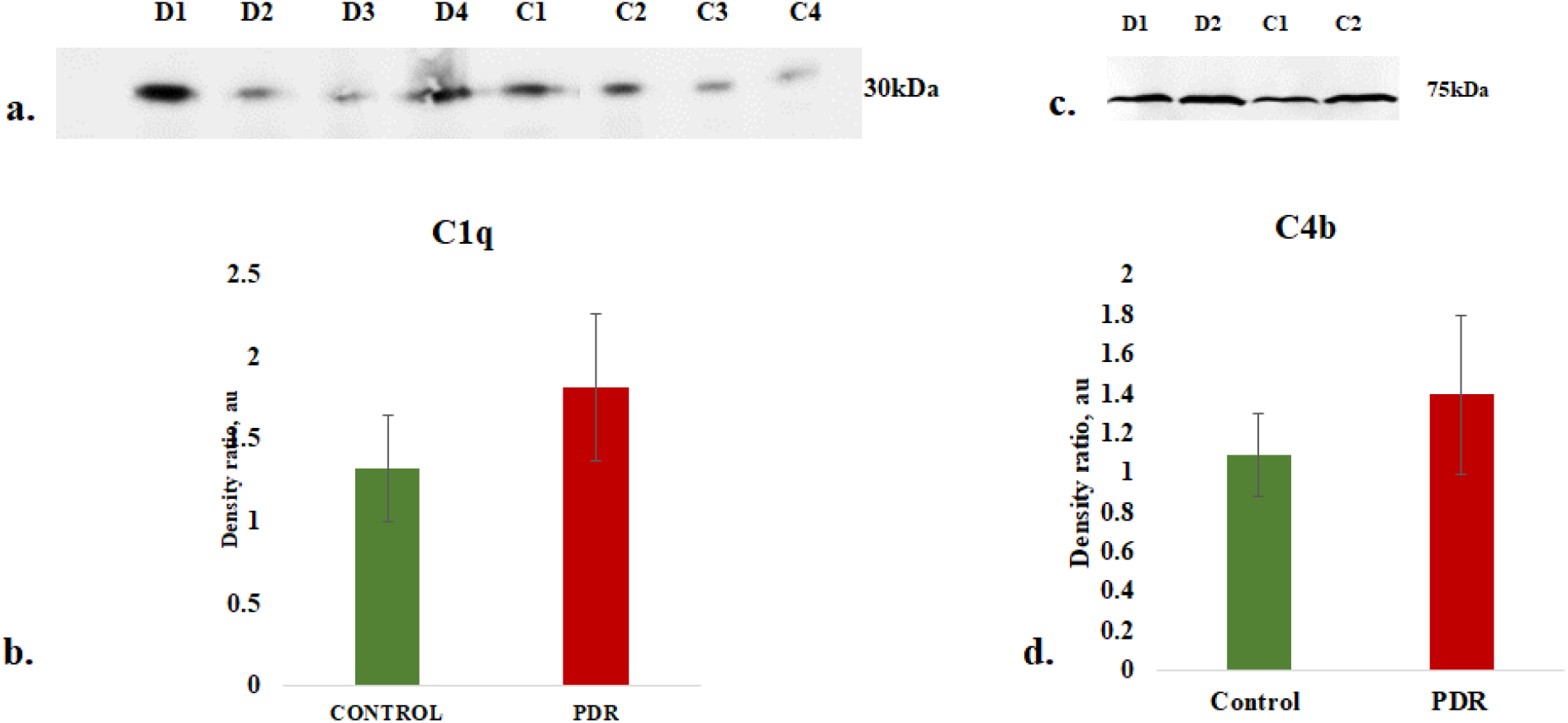
a). Representative western blots of C1q in PDR and No-DM controls, b). Quantification of C1q in PDR (n=17) and no-DM control (n=17) vitreous, *P*>*0.05,* c).Representative western blots of C1q in PDR and No-DM controls, d).Quantification of C4b in PDR (n=8), no-DM control (n=8) vitreous, *P*>*0.05* (not significant), Data represented as Mean ±SEM, C-control vitreous, D-PDR vitreous.

### 3. Contribution of alternative pathway of complement activation in diabetic retinopathy

Western blotting of CFB in vitreous samples from PDR and controls identified a band corresponding toBb fragment of CFB of molecular weight 48-50kDa (Fig 4a). Densitometry identified a significant downregulation of Bb in the PDR vitreous compared to the controls (PDR: 0.97±0.15, Controls: 1.89±0.38, *p*\*<*0.03*) (Fig 4b), indicating more bound form of Bb of factor B in the PDR vitreous for the generation of C3 convertase to activate alternative complement pathway.

**Fig 4.**
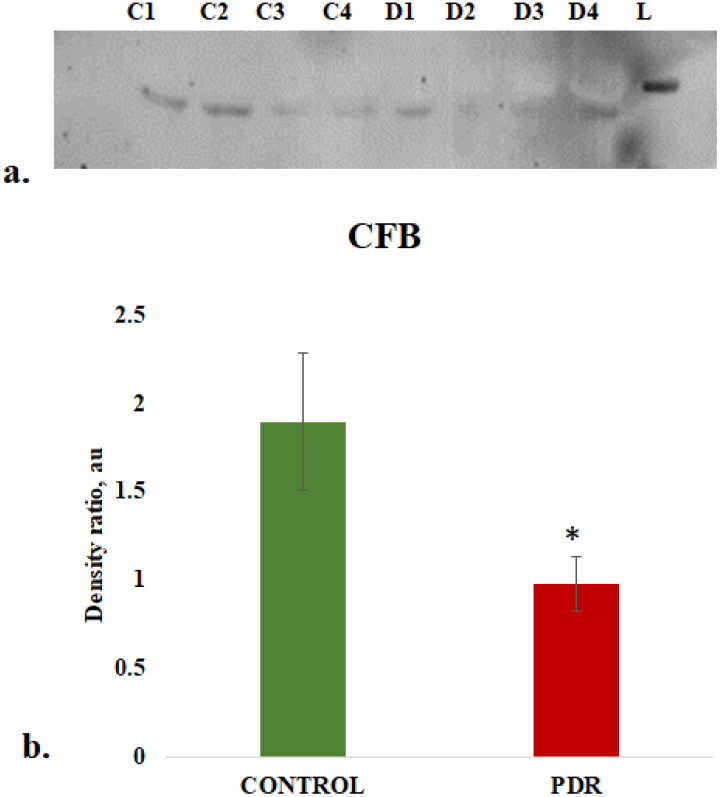
Representative western blot of CFB in PDR and No-DM controls and quantification of Bb in PDR (n=22), no-DM control (n=22) vitreous, *p*<0.05*,Data represented as Mean ±SEM,D: PDR, C: controls, L-Protein ladder

### 4. Assessment of regulation of alternative pathway of complement by complement factor H (CFH)

CFH is an important regulator of alternative pathway of complement, western blotting of CFH identified a sharp 150kDa band in vitreous samples and it was found to be significantly upregulated in PDR compared to control vitreous (Control: 0.96±0.172, PDR: 3.68±0.66, *p*\*\*<*0.0004*) (fig 5a and 5c). In order to identify whether serum infiltration is contributing to the increased level of CFH in PDR vitreous, CFH levels were measured in serum samples of no-DM, NPDR and PDR (fig 5b) and found to be downregulated for PDR as compared to NPDR and control (PDR: no-DM: 0.78±0.12, *p*>*0.05*,PDR: NPDR: 0.66±0.07, *p*>*0.05,* NPDR: no-DM*-* 1.205±0.2, *p*>*0.05*) (fig 5d).Thus, the increased level of CFH in PDR vitreous does not to be contributed by the serum infiltration and could represent a localized change in the retina mediated by one of the retinal cell types.

**Fig 5.**
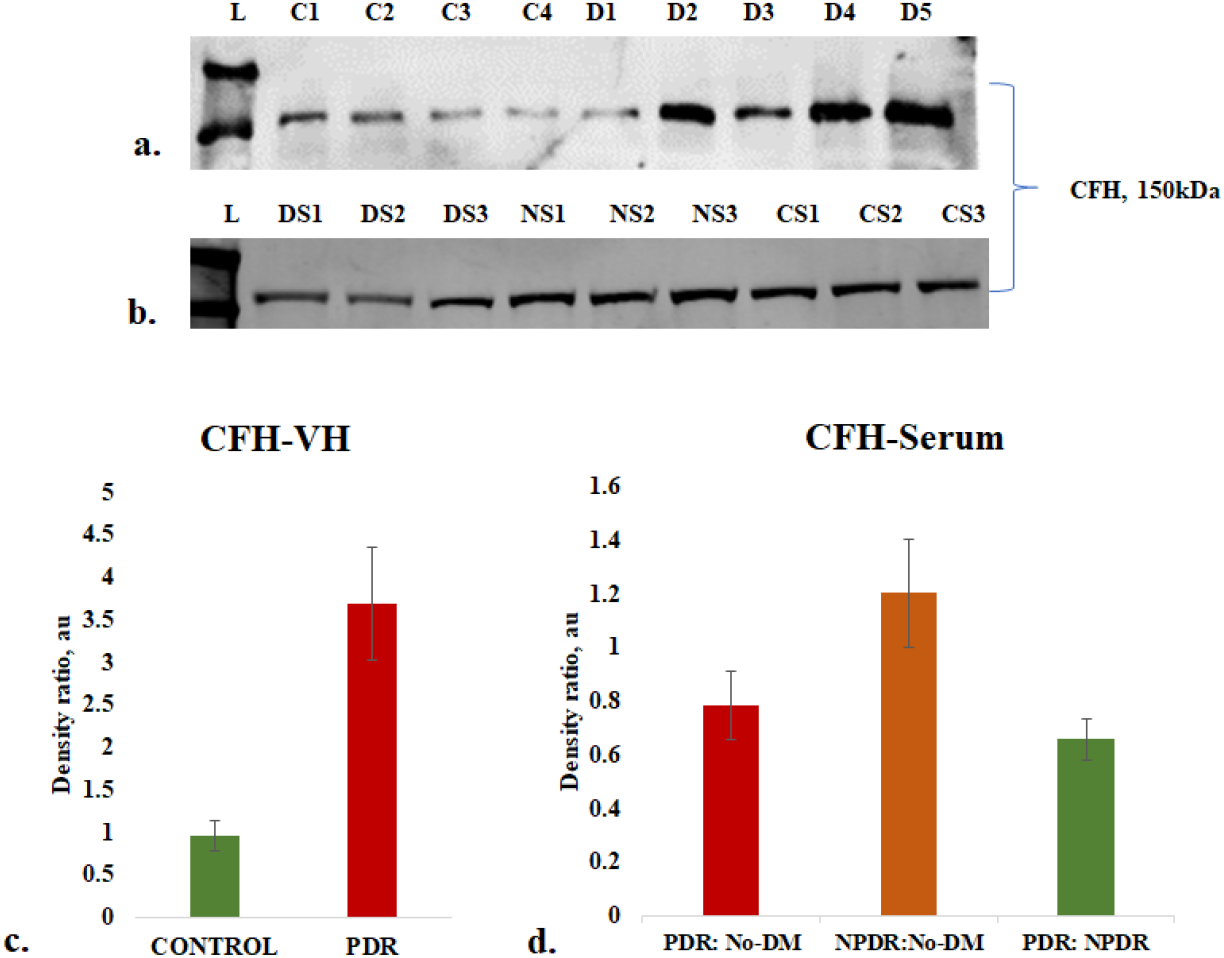
(a). Representative western blot of CFH in PDR and No-DM controls, (b). Representative western blots of CFH (150kDa) in PDR, NPDR and no-DM control serum. (c). Quantification of CFH in PDR (n=31), no-DM control (n=31) vitreous, *p**<0.0004*. (d) Quantification of CFH (150kDa) in PDR (n=12), NPDR (n=12) and no-DM control (n=12) serum, *p*>*0.05* (not significant), Data represented as Mean ±SEM, D: PDR vitreous, C: control vitreous, L-Protein ladder, DS: PDR serum, NS: NPDR and CS-Control serum, VH-Vitreous Humor.

### 5. Validation of complement activation and CFH upregulation by immunohistochemistry using diabetic and non-diabetic cadaveric retinal tissues

Diabetic and non-diabetic donor retinas were collected from cadaveric donors and performed immunohistochemistry to validate complement activation and Factor H upregulation in the retina. The retinal tissues were also stained with markers of glial activation such as CD11b for activated microglia and glial fibrillary acidic protein (GFAP) for macroglial population of the retina. Level of CXCR4 were also evaluated in the retinal tissues. The IHC results clearly demonstrated an intense staining of C3 in all the retinal layers of DM retina. GFAP was found to be present in the inner retinal layers in both DM and control retinas, however, the expression of GFAP in diabetic retina was found to be slightly higher than that of control retinas (fig 6a and 6b), suggesting the onset of gliosis in DM retinas. Increased expression of C3 and CD11b were observed in DM retina, while control retina had a relatively lower expression of these proteins (Fig.7a and 7b).

**Fig 6.**
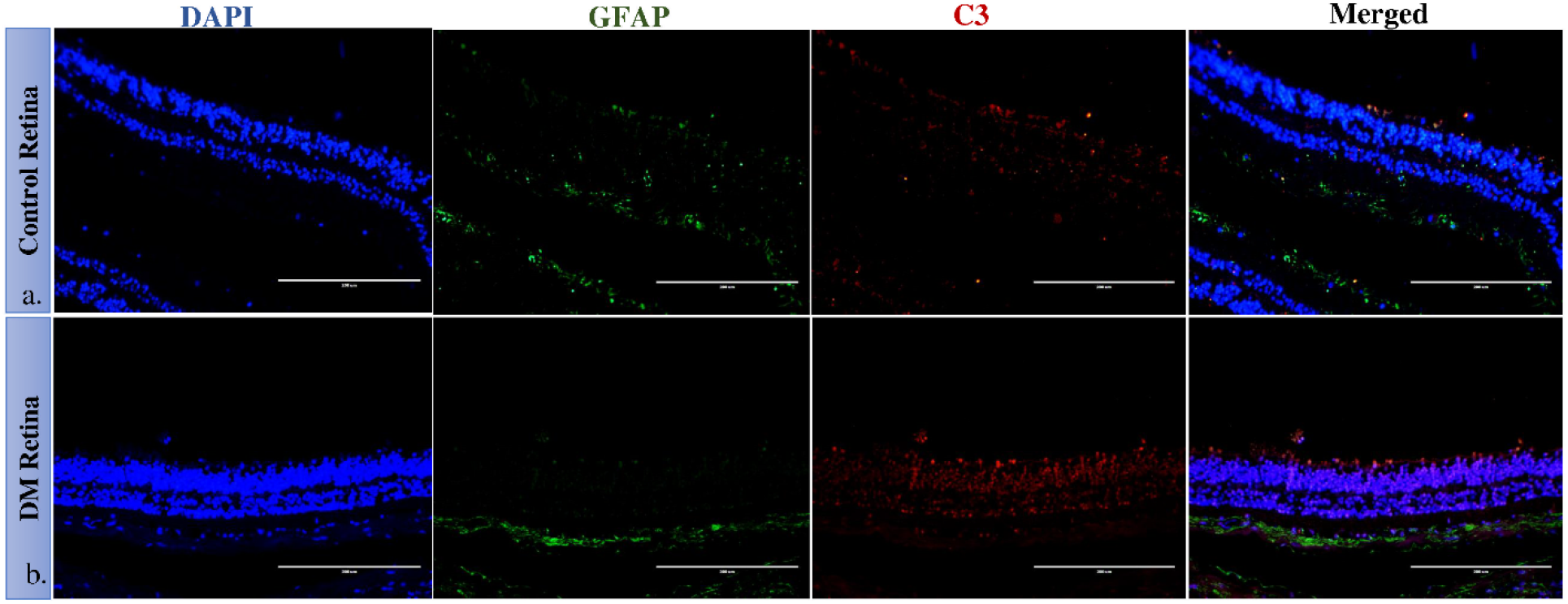
Representative immunofluorescence and localization of C3 and GFAP in retinal tissues obtained from control and diabetic retina, magnification 20X. GFAP was found to be expressed in the inner layers of the retina. C3 expression was observed in all retinal layers and intense in DM retina.

Further similar to the C3, an intense staining of CFH and its co-localized expression with CD11b positive cells in the retinal layers were observed in DM retina(Fig. 8a and 8b). This suggests that upregulation of CFH in the diabetic retina could be a feedback mechanism of excess complement activation by microglial cells. Further, the IHC results identified distribution of CXCR4+ co-localizing with CD11b positive cells in DM retina, whereas in control retinas, the CXCR4 staining was negligibly low (Fig. 9), suggestive of microglial activation and its enhanced chemotaxis under the diabetic stress.

**Fig 7.**
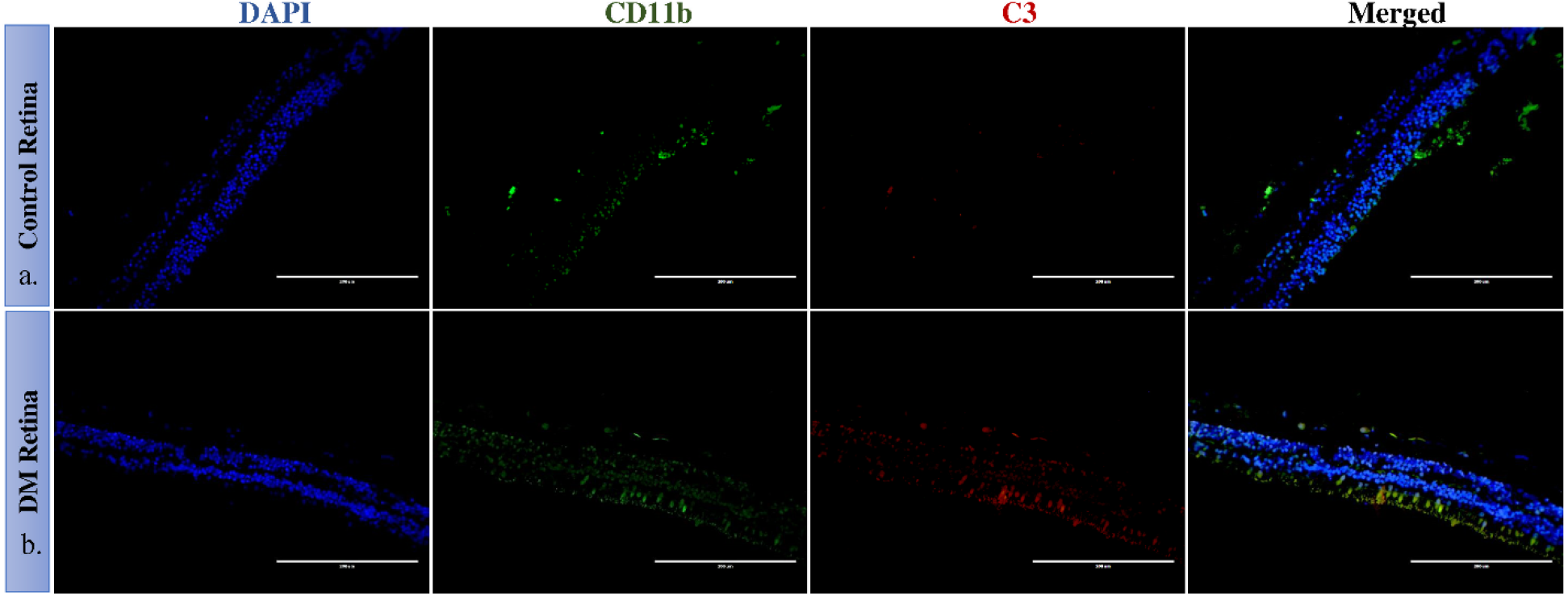
Representative immunofluorescence of C3 and CD11b in retinal tissues obtained from control and diabetic retina, magnification 20X. CD11b cells were found to be present in the inner nuclear layers in control and DM retina. In DM retina, an obvious increase in C3 expression and CD11b was observed.

**Fig 8.**
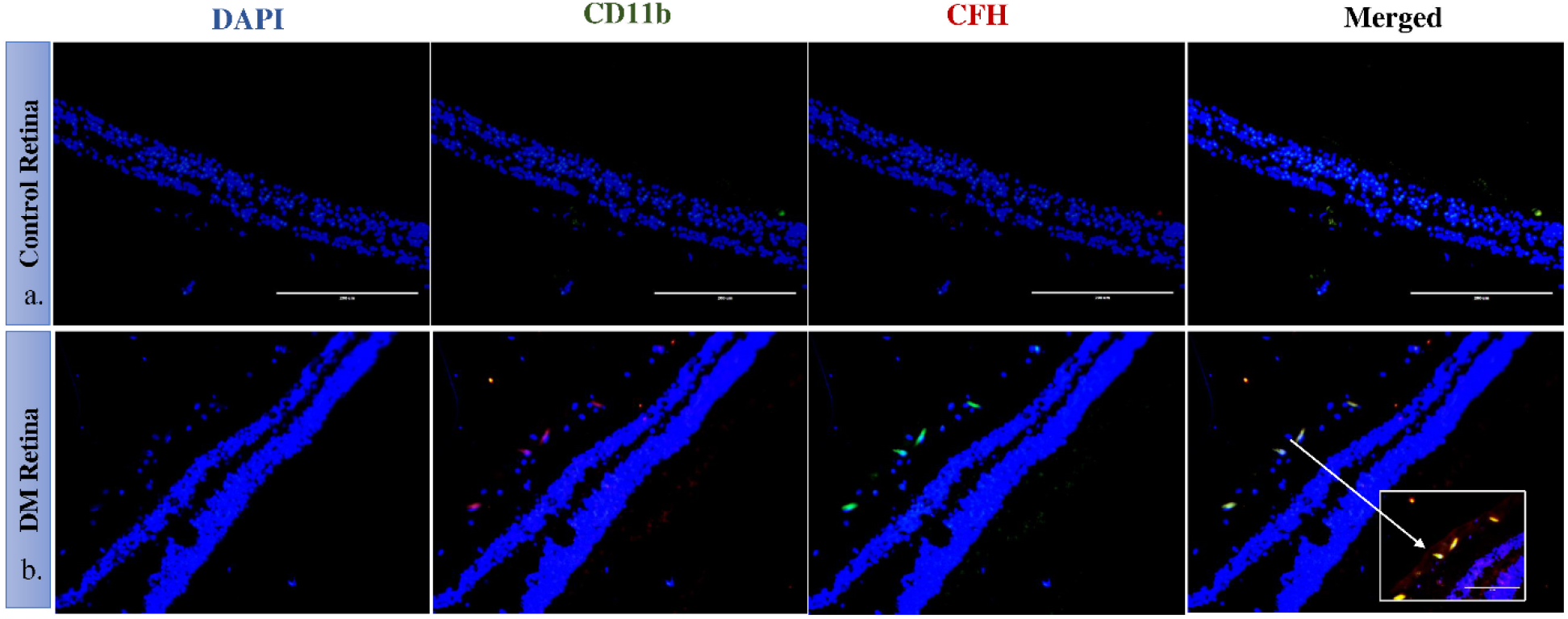
Representative immunofluorescence of and localization of CFH and CD11b in retinal tissues obtained from control and diabetic retina, magnification 20X. Co-localized expression of CFH and CD11b is highlighted in 40X magnification in the panel B

**Fig 9:**
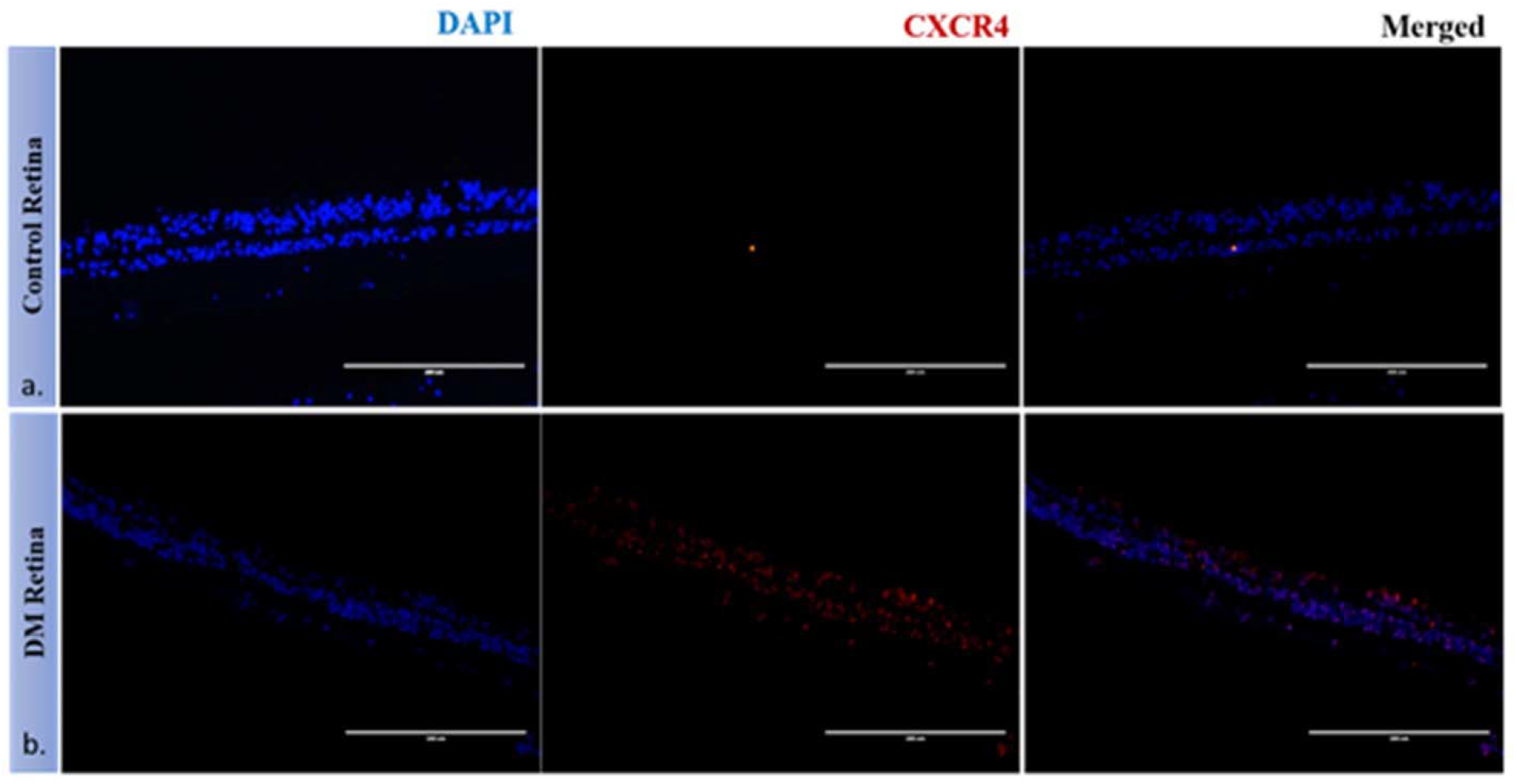
Representative immunofluorescence of CXCR4 in retinal tissues obtained from control and diabetic retina, magnification 20X. A clear invasion of CXCR4 cells is seen

### 6. Assessment of microglial activation by analysing the level of matrix metalloproteinases (MMPs)

Microglia are the major source of gelatinolytic MMPs such as MMP9 and MMP2 in the retina. In order to evaluate the enzymatic activity of MMPs in the PDR vitreous gelatin zymography was done and compared MMP activity with vitreous samples from controls. The zymography results identified a clear gelatinolytic band in the PDR and control vitreous at 82-85kDa molecular weight, which corresponds to active MMP9. This band was found to be more pronounced in PDRs than control vitreous, indicating more gelatinolytic activity and active MMP9 in the PDR cases as compared to controls (fig 10a).

**Fig 10:**
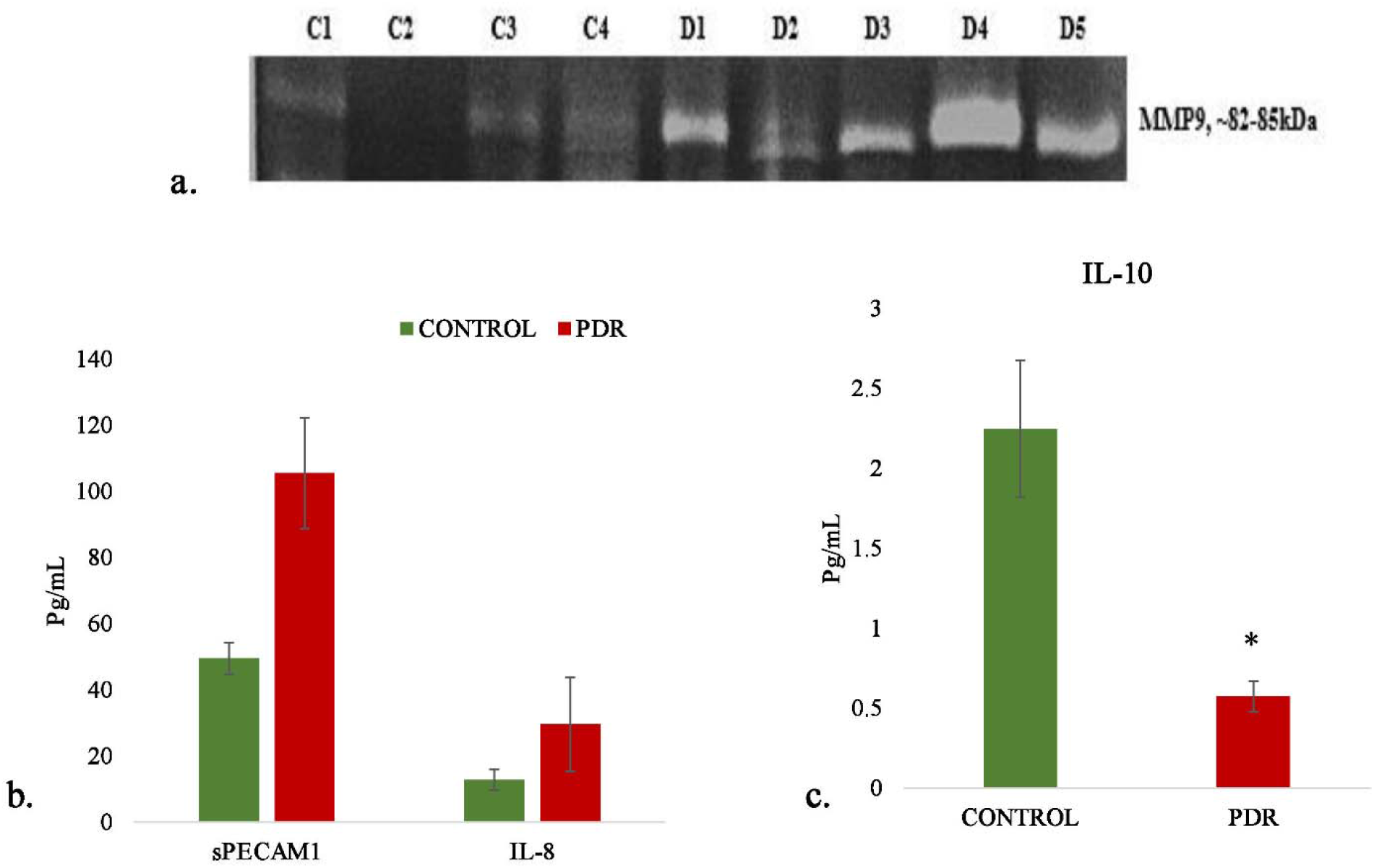
a. Representative Gelatin zymography of vitreous samples from PDR (n=10) as compared to controls (n=10) (C-control, D-PDR),(b). Quantitative estimation of sPECAMIL-8and (c) IL-10 in PDR vs Control vitreous by multiplex ELISA, \**p<0.05,* PDR=8 and Control=8

### 7. Analysis of inflammation and microglial activation in vitreous samples by analyzing the level of sPECAM, IL-8 and IL-10

The level of inflammation in vitreous samples were analysed by the quantitative estimation of pro-inflammatory markers such as sPECAM, IL-8 and also the estimation of anti-inflammatory marker IL-10. The level of sPECAM and IL-8 were found to be higher in PDR vitreous as compared to the control vitreous. (sPECAM-Control (n=8): 49.54±4.76, PDR:105.45±16.69, *p*\*<*0.01*, IL-8: Control (n=8), 12.79±3.13, PDR (n=8): 29.47±14.14, *p*>*0.05*, non-significant). In contrast to this, the level of anti-inflammatory cytokine IL-10 was found to be significantly downregulated in PDR vitreous as compared to control (Control (n=8): 2.24±0.42, PDR (n=8): 0.57±0.09 *p*\*\*<*0.001*) (Fig 10 b and 10c).

### 8. Analysis of complement and angiogenic genes by quantitative real time PCR

Blood samples were collected and RNA was isolated from PDR, NPDR and control subjects and quantitative real time PCR was performed for the genes such as *TGF* β1, *THSB1, CXCR4, VEGF, C3* and *CFH*. The gene expression analysis identified the expression level of these genes as, *TGF* β-NPDR (n=20): 0.92 ± 0.22 (*p*>*0.05, N.S*), PDR (n=20) 0.73 ±0.08 (*p*\*<*0.05*), *THSB1*-NPDR (n=20): 0.499±0.15 (*p*\*\*<*0.004*), PDR (n=20): 0.495 ± 0.06 (*p*\*\*<*0.008), VEGF*-NPDR (n=20): 1.43± 0.21 (*p*>*0.05, N.S*), PDR (n=20): 1.53 ± 0.07 (*p*\*<*0.02*), *C3*-NPDR (n=20): 1.17 ±0.173 (*p*>*0.05, N.S*), PDR (n=20): 1.45±0.14 (*p*\*<*0.01*), CFH – NPDR (n=20): 0.628 ± 0.13 (*p*\*<*0.0001*), PDR (n=20), 0.45± 0.13 (*p*\*<*0.0003*) and CXCR4-NPDR (n=20), 1.03± 0.18 (*p*>*0.05, N.S*), PDR (n=20): 1.21±0.22 (*p*>*0.05, N.S*) (Fig 11).

**Fig 11:**
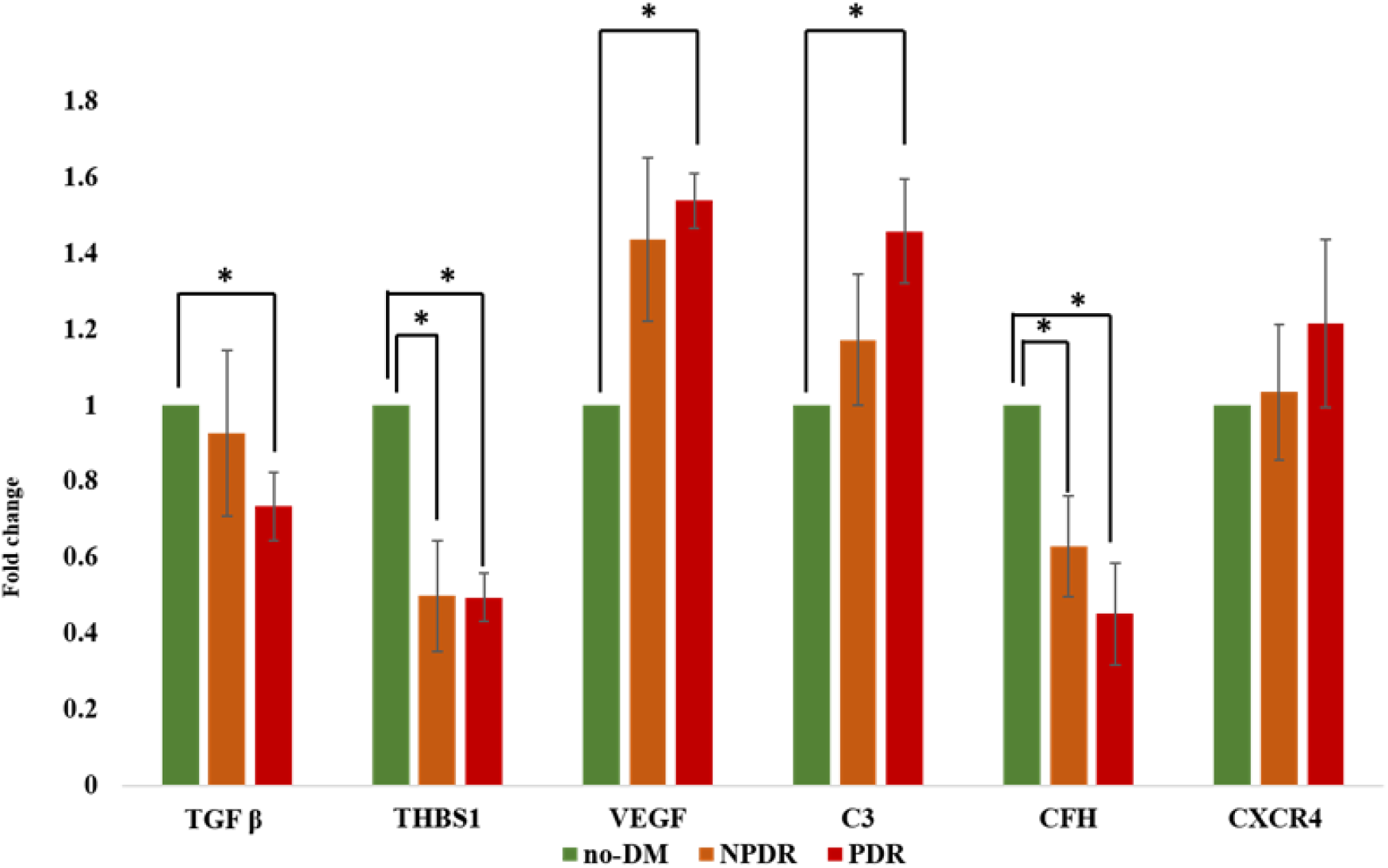
Quantitative real time PCR of complement and angiogenic genes in PDR (n=20), NPDR (n=20) compared to no-DM controls (n=20), \**p*<0.05, Date represented as Mean ± SEM

## Discussion

Diabetic retinopathy is a serious neuro-vascular complications of retina and involvement of complement pathway in DR progression is paid more attention after the identification of complement deposits in choriocapillaries of DR retina and reduced levels of complement pathway inhibitors in diabetic retina[16, 17]. A comprehensive study done by Garcia *et al*, identified a significant increase in 42kDa fragment of C3 and CFB in PDR vitreous and found it to correlate with the mRNA expression in diabetic retina [22]. Additionally, various vitreous proteome studies in DR also identified the predominant deposition of complement proteins in PDR vitreous [20, 21, 26, 27] though exactly how complements are involved in PDR pathogenesis was not explored. Thus, here for the first time, we attempted a systematic evaluation of both alternative and classical pathwayof complement activation in PDR pathogenesis by analysing proteins of these two pathways in serum as well as in vitreous humor. The major rationale behind this comparison is that, if these proteins are contributed by blood retinal barrier breakdown in advance stages of the disease. Hence, we also checked for the systemic complement activation in serum from PDR, NPDR and no-DM subjects to make sure if complement dysregulation as seen in vitreous proteome in PDR is a localized effect and not an additive effect due to serum infiltration.

C3 is the central complement protein, where all the three complement pathways converge. The activation of complement pathway causes proteolytic fragmentation of C3 and these fragments can bind to the nearby tissues and enhance inflammatory process [28]. In our study cohort, no significant increase in total C3 or any of its activated fragments was seen among PDR and NPDR serum compared to control serum, while a significant increase in total C3 was observed in PDR vitreous. Though, there was no significant differences in number of activated C3 fragments in PDR vitreous compared to control, suggesting significant complement activation in control vitreous as well. This could also be attributed to homeostatic changes in retinal microenvironment in conditions of macular hole and retinal detachment cases that serve as controls for this experiment. However, a significant upregulation of C3bα’(110kDa) fragment was noted in the PDR vitreous. The reactive C3bα’ is generated from 120kDa α-chain of C3 after the proteolytic removal of 10kDa C3a fragment and is the part of active C3b [29]. A significant increase in the level of C3bα’indicated for complement pathway activation in PDR vitreous. C1q and C4b are the proteins specific for classical pathway of complement activation and both these proteins identified an insignificant increase of C1q in PDR and NPDR serum and C1q and C4b in PDR vitreous. This suggested that classical complement pathway might not be playing asignificant role in PDR pathogenesis.

Further, we have looked into the alternative complement pathway by selecting CFB, a protein specific for alternative pathway that is required for the formation of C3 convertase (C3bBb) and activates the alternative pathway of complement[30]. Fragment Bb of factor B was found to be significantly downregulated in PDR vitreous which could be due to the formation of more C3bBb in PDR vitreous. Next, we tested the levels of CFH, one of the major regulators of alternative complement pathway. CFH regulate alternative complement pathway in multiple steps such as competing with FB for C3b binding, also act as a co-factor for factor I to degrade C3b to C3bi [31]. We have expected a low level of CFH in PDR vitreous after observing the downregulation of free Bb in the PDR vitreous. But surprisingly, we found a significant upregulation of *CFH* gene expression in PDR vitreous. Several studies have shown that one of the binding sites of CFH for C3b is located in the C3bα’ region[32], and thus quite possibly the significant upregulation of CFH could be a feed-back mechanism for maintaining the level of C3bα’ in the PDR vitreous. Further the serum CFH levels on the other hand, showed a down-regulation of this protein in PDR cases compared to NPDR and control, suggesting thereby that the upregulation of CFH as seen in vitreous is a localized phenomenon.

Further immunohistochemistry experiments confirmed the CFH and C3 upregulation as seen in PDR vitreous, in the retinal tissues collected from diabetic donor eyes as compared to the control non diabeticdonor’sretina. Similar to the vitreous data, we found a concurrent deposition of C3 and CFH in the diabetic retina. To test further if this CFH and C3 overexpression as seen in retinal tissues from diabetic donor by microglial cells as seen in AMD and ROP conditions, we performed co-localization of CFH and C3 proteins with activated microglial specific marker CD11b. A significant amount of CFH expression was seen to localize with microglial cell that were present in the inner nuclear layers. We also noticed significant gliosis in the diabetic tissues as evident from the upregulation of GFAP protein in diabetic retina. However, it was microglial cells and not the macroglia that seemed to be the major source of complement proteins in diabetic retina. Further confirmation of the microglial activation as seen in PDR cases, was evident from the significant down regulation of IL-10 in the PDR vitreous samples while IL-8 was upregulated (though not statistically significant). IL-8 is a proinflammatory and IL-10 is an anti-inflammatory cytokine secreted mainly by M1 and M2 microglia [33]. The down regulation of IL-10 and upregulation of IL-8 further confirmed activation of proinflammatory M1 phenotypes in PDR vitreous. Microglial once activated are known to secrete the matrix metalloproteinases and this too was confirmed using. zymography that showed significantly increased level of MMP9 secretion in PDR vitreous[34]. This further validated microglial activation in PDR vitreous. Alongside an increase in proinflammatory markers such as Il-8 and MMP9, a significant upregulation of sPECAM in the PDR vitreous, also indicated excessive inflammation in the PDR vitreous. This also suggests for an inflammatory environment in the PDR retina that could be a driving force for microglial activation. Microglial cells being the resident cells gets activated and move up from deep RGC layers towards photoreceptors. Microglial activation was further evaluated in the diabetic retina using CXCR4, which is a chemokine receptorand known to be involved in astroglial activation and microglial signaling. Our IHC results identified an increased expression of these receptors in the inner retinal layers, suggesting microglial activation and its migration in diabetic retina.

Next, we correlated the increase in microglial cells with the mRNA expression of C3, CFH, THSB1, VEGF, TGFβ and CXCR4 in PBMNC isolated from PDR, NPDR and no-DM cases. Our gene expression data showed upregulation of C3 in PDR and NPDR cases and a concurrent downregulation of CFH. This was found to be consistent with the serum level C3 and CFH expression, where we found insignificant upregulation of C3 and downregulation of CFH in PDR and NPDR cases. Further the level of *THSB1*, a potent anti-angiogenic gene and also the activator of *TGF*β*1*[35] were evaluated and we found a significant downregulation of THSB1 in PDR and NPDR cases, simultaneously pro-inflammatory gene *TGF* β1 also found to be downregulated. Our study also confirmed that THSB1 is a negative fluid phase regulator of alternative pathway of complement activation[36].and therefore the downregulation of THSB1 further correlated with the complement pathway activation in DR.

In conclusion, our study provided a systematic analysis of classical and alternative complement pathway activation in PDR pathogenesis. We for the first time showed a significant upregulation of 110kDa C3bα’and concurrent increase of CFH in PDR vitreous, and that this upregulation of complement cascade was localized to retina and not contributed by the blood retinal barrier breakdown which is common sight in advance PDR cases. Further, a correlation of increased Complement factor H expression and downregulation of Complement factor B with genetic changes in these genes could underscore the contribution of the alternate complement pathway in DR pathogenesis Lastly, our study suggested that activated microglia are the main source of alternative pathway of complement activation in PDR. In future, targeting microglial mediated complement activation could pave way for an effective therapeutic management of DR by the reducing underlying inflammation and abnormal angiogenesis.

## References

1. Chen M, Xu H. Parainflammation, chronic inflammation, and age-related macular degeneration. Journal of leukocyte biology.2015; 98:713–725.

2. Mukai R, Okunuki Y, Husain D, Kim CB, Lambris JD, Connor KM. The Complement System Is Critical in Maintaining Retinal Integrity during Aging. Frontiers in aging neuroscience.2018; 10:15.

3. McGeer PL, McGeer EG. Inflammation and the degenerative diseases of aging. Annals of the New York Academy of Sciences.2004; 1035:104–116.

4. Strey CW, Markiewski M, Mastellos D, Tudoran R, Spruce LA, Greenbaum LE, Lambris JD. The proinflammatory mediators C3a and C5a are essential for liver regeneration. The Journal of experimental medicine.2003; 198:913–923.

5. Stevens B, Allen NJ, Vazquez LE, Howell GR, Christopherson KS, Nouri N, Micheva KD, Mehalow AK, Huberman AD, Stafford B, et al. The classical complement cascade mediates CNS synapse elimination. Cell.2007; 131:1164–1178.

6. Langer HF, Chung KJ, Orlova VV, Choi EY, Kaul S, Kruhlak MJ, Alatsatianos M, DeAngelis RA, Roche PA, Magotti P, et al. Complement-mediated inhibition of neovascularization reveals a point of convergence between innate immunity and angiogenesis. Blood.2010; 116:4395–4403.

7. Rajappa M, Saxena P, Kaur J. Ocular angiogenesis: mechanisms and recent advances in therapy. Advances in clinical chemistry.2010; 50:103–121.

8. Bora PS, Sohn JH, Cruz JM, Jha P, Nishihori H, Wang Y, Kaliappan S, Kaplan HJ, Bora NS. Role of complement and complement membrane attack complex in laser-induced choroidal neovascularization. Journal of immunology.2005; 174:491–497.

9. Rathi S, Jalali S, Patnaik S, Shahulhameed S, Musada GR, Balakrishnan D, Rani PK, Kekunnaya R, Chhablani PP, Swain S, et al. Abnormal Complement Activation and Inflammation in the Pathogenesis of Retinopathy of Prematurity. Frontiers in immunology.2017; 8:1868.

10. Cao S, Wang JC, Gao J, Wong M, To E, White VA, Cui JZ, Matsubara JA. CFH Y402H polymorphism and the complement activation product C5a: effects on NF-kappaB activation and inflammasome gene regulation. The British journal of ophthalmology.2016; 100:713–718.

11. Edwards AO, Ritter R, 3rd, Abel KJ, Manning A, Panhuysen C, Farrer LA. Complement factor H polymorphism and age-related macular degeneration. Science.2005; 308:421–424.

12. Mantel I, Ambresin A, Moetteli L, Droz I, Roduit R, Munier FL, Schorderet DF. Complement factor B polymorphism and the phenotype of early age-related macular degeneration. Ophthalmic genetics.2014; 35:12–17.

13. Tanaka K, Sonoo H, Kurebayashi J, Nomura T, Ohkubo S, Yamamoto Y, Yamamoto S. Inhibition of infiltration and angiogenesis by thrombospondin-1 in papillary thyroid carcinoma. Clinical cancer research : an official journal of the American Association for Cancer Research.2002; 8:1125–1131.

14. Zamiri P, Masli S, Kitaichi N, Taylor AW, Streilein JW. Thrombospondin plays a vital role in the immune privilege of the eye. Investigative ophthalmology & visual science.2005; 46:908–919.

15. Yau JW, Rogers SL, Kawasaki R, Lamoureux EL, Kowalski JW, Bek T, Chen SJ, Dekker JM, Fletcher A, Grauslund J, et al. Global prevalence and major risk factors of diabetic retinopathy. Diabetes care.2012; 35:556–564.

16. Gerl VB, Bohl J, Pitz S, Stoffelns B, Pfeiffer N, Bhakdi S. Extensive deposits of complement C3d and C5b-9 in the choriocapillaris of eyes of patients with diabetic retinopathy. Investigative ophthalmology & visual science.2002; 43:1104–1108.

17. Zhang J, Gerhardinger C, Lorenzi M. Early complement activation and decreased levels of glycosylphosphatidylinositol-anchored complement inhibitors in human and experimental diabetic retinopathy. Diabetes.2002; 51:3499–3504.

18. Fujita T, Hemmi S, Kajiwara M, Yabuki M, Fuke Y, Satomura A, Soma M. Complement-mediated chronic inflammation is associated with diabetic microvascular complication. Diabetes/metabolism research and reviews.2013; 29:220–226.

19. Gao BB, Chen X, Timothy N, Aiello LP, Feener EP. Characterization of the vitreous proteome in diabetes without diabetic retinopathy and diabetes with proliferative diabetic retinopathy. Journal of proteome research.2008; 7:2516–2525.

20. Wang H, Feng L, Hu J, Xie C, Wang F. Differentiating vitreous proteomes in proliferative diabetic retinopathy using high-performance liquid chromatography coupled to tandem mass spectrometry. Experimental eye research.2013; 108:110–119.

21. Li J, Lu Q, Lu P. Quantitative proteomics analysis of vitreous body from type 2 diabetic patients with proliferative diabetic retinopathy. BMC ophthalmology.2018; 18:151.

22. Garcia-Ramirez M, Canals F, Hernandez C, Colome N, Ferrer C, Carrasco E, Garcia-Arumi J, Simo R. Proteomic analysis of human vitreous fluid by fluorescence-based difference gel electrophoresis (DIGE): a new strategy for identifying potential candidates in the pathogenesis of proliferative diabetic retinopathy. Diabetologia.2007; 50:1294–1303.

23. Yang MM, Wang J, Ren H, Sun YD, Fan JJ, Teng Y, Li YB. Genetic Investigation of Complement Pathway Genes in Type 2 Diabetic Retinopathy: An Inflammatory Perspective. Mediators of inflammation.2016; 2016:1313027.

24. Wang J, Yang MM, Li YB, Liu GD, Teng Y, Liu XM. Association of CFH and CFB gene polymorphisms with retinopathy in type 2 diabetic patients. Mediators of inflammation.2013; 2013:748435.

25. Toth M, Fridman R. Assessment of Gelatinases (MMP-2 and MMP-9 by Gelatin Zymography. Methods in molecular medicine.2001; 57:163–174.

26. Hernandez C, Garcia-Ramirez M, Colome N, Corraliza L, Garcia-Pascual L, Casado J, Canals F, Simo R. Identification of new pathogenic candidates for diabetic macular edema using fluorescence-based difference gel electrophoresis analysis. Diabetes/metabolism research and reviews.2013; 29:499–506.

27. Loukovaara S, Nurkkala H, Tamene F, Gucciardo E, Liu X, Repo P, Lehti K, Varjosalo M. Quantitative Proteomics Analysis of Vitreous Humor from Diabetic Retinopathy Patients. Journal of proteome research.2015; 14:5131–5143.

28. Nishida N, Walz T, Springer TA. Structural transitions of complement component C3 and its activation products. Proceedings of the National Academy of Sciences of the United States of America.2006; 103:19737–19742.

29. Clay CD, Soni S, Gunn JS, Schlesinger LS. Evasion of complement-mediated lysis and complement C3 deposition are regulated by Francisella tularensis lipopolysaccharide O antigen. Journal of immunology.2008; 181:5568–5578.

30. Noris M, Remuzzi G. Overview of complement activation and regulation. Seminars in nephrology.2013; 33:479–492.

31. Schmidt CQ, Herbert AP, Hocking HG, Uhrin D, Barlow PN. Translational mini-review series on complement factor H: structural and functional correlations for factor H. Clinical and experimental immunology.2008; 151:14–24.

32. Jokiranta TS, Hellwage J, Koistinen V, Zipfel PF, Meri S. Each of the three binding sites on complement factor H interacts with a distinct site on C3b. The Journal of biological chemistry.2000; 275:27657–27662.

33. Martinez FO, Gordon S. The M1 and M2 paradigm of macrophage activation: time for reassessment. F1000prime reports.2014; 6:13.

34. del Zoppo GJ, Frankowski H, Gu YH, Osada T, Kanazawa M, Milner R, Wang X, Hosomi N, Mabuchi T, Koziol JA. Microglial cell activation is a source of metalloproteinase generation during hemorrhagic transformation. Journal of cerebral blood flow and metabolism : official journal of the International Society of Cerebral Blood Flow and Metabolism.2012; 32:919–932.

35. Crawford SE, Stellmach V, Murphy-Ullrich JE, Ribeiro SM, Lawler J, Hynes RO, Boivin GP, Bouck N. Thrombospondin-1 is a major activator of TGF-beta1 in vivo. Cell.1998; 93:1159–1170.

36. http://hdl.handle.net/2381/40955

