## Supplementary data for "A systematic investigation on the involvement of complement pathway in diabetic retinopathy"

**Inderjeet Kaur**

Prof Brien Holden Eye Research Centre, LV Prasad Eye Institute, Hyderabad, India,

Supplementary Table S1: Demographics of study subjects used for vitreous protein analysis

|  | Age | Gender | Duration of DM |
| --- | --- | --- | --- |
| Control Vitreous | 55.4±1.02 | F, n= 60, M, n =40 | Nil |
| PDR vitreous | 56.17±0.79 | F, n = 45, M, n =55 | 15.64±0.83 |

Supplementary table S2: Detailed demographics of study subjects used for serum protein analysis and mRNA expression analysis by qPCR.

|  | Age | Gender | Duration of DM |
| --- | --- | --- | --- |
| No-DM | 65.8±1.03 | F, n=16, M, n =22 | Nil |
| NPDR | 59.83±1.32 | F, n = 14, M, n =24 | 12.88±1.4 |
| PDR | 53.86±1.61 | F, n = 15, M, n =23 | 15.05±0.9 |

Supplementary table S3: Details of antibodies used for western blotting and their dilutions.

| Antibody list | Conc. of protein used | Dilution | % of gel and condition used |
| --- | --- | --- | --- |
| Ms C3, Santacruz-sc-28294 | 15 μg | 1:300 | 7.5% SDS PAGE, non-reducing condition |
| Ms C1q, Abcam-ab71089 | 30μg | 1:1000 | 12% SDS PGAE, reducing condition |
| Ms C4b, Santa Cruz, sc-74524 | 25μg | 1:200 | 10%SDS PAGE, non-reducing condition |
| Rb CFB, Abcam, ab72658 | 25μg | 1:500 | 10%SDS PAGE, non-reducing condition |
| Ms CFH, Santacruz, sc-166613 | 30 μg | 1:200 | 7.5%SDS PAGE, reducing condition |
| Anti -Ms. 680RD, Licor-cat. no. 926-68070 | - | 1:75,00 | - |
| Anti -Rb 800CW, Licor, cat.no.926-32211 | -- | 1:10,000 | -- |

Supplementary table S4 Primer sequence and TaqMan assays used for qRT PCR

| Gene | Forward primer | Reverse primer | Ref. |
| --- | --- | --- | --- |
| CXCR4 | AGCATGACGGACAAGTACAGG | GATGAAGTCGGGAATAGTCAGC | ^1^ |
| Vegf 165 | ATCTTCAAGCCATCCTGTGTGC | CAAGGCCCACAGGGATTTTC | ^2^ |
| CFH | TACTGGCTGGATACCTGCTC | CCTGACGGAGTCTCAAAATG | ^3^ |
| β Actin | TCTACAATGAGCTGCGTGTG | GGTGAGGATCTTCATGAGGT | ^4^ |

Supplementary Table S5: Complement proteins and their corresponding number of peptides identified in no-DM, DM and PDR vitreous by LC-MS/MS

| Complement proteins | No-DM peptide (Mean ± SEM) (n=3) | DM peptide  (Mean ± SEM) (n=3) | PDR peptide  (Mean ± SEM) (n=3) |
| --- | --- | --- | --- |
| C9 | 55±2.5 | 37.6±4.3 | 42±9.1 |
| CRP | 9±1 | 5±0 | 5.6±0.8 |
| C1QB | 12.3±1.8 | 12.3±1.7 | 12±1.1 |
| C1QC | 11.33±0.9 | 12±0.5 | 9±1 |
| C1S | 26±5.8 | 25±7.7 | 18.3±3.3 |
| C2 | 7.3±1.2 | 6±0.57 | 5±0.5 |
| C3 | 532.6±39.8 | 525.6±56 | 523.6±86.7 |
| C4A | 19.6±4.5 | 18.6±4.4 | 17±3.5 |
| C4BPA | Nil | Nil | 3.3±0.88 |
| C5 | 53.3±4.3 | 37.3±5.17 | 44.3±7.3 |
| CLU | 222.6±7.5 | 221±24.1 | 174.6±27.5 |
| C1RL | 10±0.58 | 7±0.5 | 6±2.08 |
| C7 | 23.3±2 | 23.6±4.3 | 19.6±4.4 |
| C6 | 23±0.5 | 16.3±2.1 | 20.3±2.4 |
| C8A | 16.3±2.6 | 19.3±3.2 | 14±4.35 |
| C8G | 17.3±1.8 | 17±2.5 | 14.3±1.85 |
| CFH | 52±2.5 | 49±7.8 | 57±5.7 |

1.	Vaday GG, Hua SB, Peehl DM, et al. CXCR4 and CXCL12 (SDF-1) in prostate cancer: inhibitory effects of human single chain Fv antibodies. *Clinical cancer research : an official journal of the American Association for Cancer Research* 2004;10:5630-5639.

2.	Medford AR, Douglas SK, Godinho SI, et al. Vascular Endothelial Growth Factor (VEGF) isoform expression and activity in human and murine lung injury. *Respiratory research* 2009;10:27.

3.	Zhang Y, Huang Q, Tang M, Zhang J, Fan W. Complement Factor H Expressed by Retinal Pigment Epithelium Cells Can Suppress Neovascularization of Human Umbilical Vein Endothelial Cells: An in vitro Study. *PloS one* 2015;10:e0129945.

4.	Horvatinovich JM, Sparks SD, Borowitz MJ. Detection of terminal deoxynucleotidyl transferase by flow cytometry: a three color method. *Cytometry* 1994;18:228-230.
